# Three-way relationships between gut microbiota, helminth assemblages and bacterial infections in wild rodent populations

**DOI:** 10.1101/2022.05.23.493084

**Authors:** Marie Bouilloud, Maxime Galan, Adélaïde Dubois, Christophe Diagne, Philippe Marianneau, Benjamin Roche, Nathalie Charbonnel

**Affiliations:** CBGP, INRAE, CIRAD, Institut Agro, IRD, University of Montpellier, Montpellier, France; MIVEGEC, IRD, CNRS, University of Montpellier, Montpellier, France; CBGP, IRD, INRAE, CIRAD, Institut Agro, University of Montpellier, Montpellier, France; INRAE, Lyon, France

**Author notes:** Equal contribution of the coauthors (Benjamin Roche, Nathalie Charbonnel).

**Keywords:** bank voles, co-infections, interactions, microbial community ecology, zoonoses

## Abstract

**Background:** Despite its central role in host fitness, the gut microbiota may differ greatly between individuals. This variability is often mediated by environmental or host factors such as diet, genetics, and infections. Recently, a particular attention has been given to the interactions between gut bacteriota and helminths, as these latter could affect host susceptibility to other infections. Further studies are still required to better understand the three-way interactions between gut bacteriota, helminths and other parasites, especially because previous findings have been very variable, even for comparable host-parasite systems.

**Methods:** In our study, we used the V4 region of the 16S rRNA gene to assess the variability of gut bacteriota diversity and composition in wild populations of a small mammal, the bank vole *Myodes glareolus*. Four sites were sampled at a regional geographical scale (100 km) along a North-South transect in Eastern France. We applied analyses of community and microbial ecology to evaluate the interactions between the gut bacteriota, the gastro-intestinal helminths and the pathogenic bacteria detected in the spleen.

**Results:** We identified important variations of the gut bacteriota composition and diversity among bank voles. They were mainly explained by sampling localities and reflected the North/South sampling transect. In addition, we detected two main enterotypes, that might correspond to contrasted diets. We found geographic variations of the Firmicutes/Bacteroidetes ratio, that correlated positively with body mass index. We found positive correlations between the specific richness of the gut bacteriota and of the helminth community, as well as between the composition of these two communities, even when accounting for the influence of geographical distance. The helminths *Aonchotheca murissylvatici, Heligmosomum mixtum* and the bacteria *Bartonella* sp were the main taxa associated with the whole gut bacteriota composition. Besides, changes in relative abundance of particular gut bacteriota taxa were specifically associated with other helminths (*Mastophorus muris, Catenotaenia henttoneni, Paranoplocephala omphalodes* and *Trichuris arvicolae*) or pathogenic bacteria. Especially, infections with *Neoehrlichia mikurensis, Orientia* sp, *Rickettsia* sp and *P. omphalodes* were associated with lower relative abundance of the family Erysipelotrichaceae (Firmicutes), while coinfections with higher number of bacterial infections were associated with lower relative abundance of a Bacteroidales family (Bacteroidetes).

**Conclusions:** These results emphasize complex interlinkages between gut bacteriota and infections in wild animal populations. They remain difficult to generalize due to the strong impact of environment on these interactions, even at regional geographical scales. Abiotic features, as well as small mammal community composition and within host parasite coinfections, should now be considered to better understand the spatial variations observed in the relationships between gut bacteriota, gastro-intestinal helminths and bacterial infections.

## Introduction

Vertebrate gut microbiota plays key roles in host fitness through functions including nutrient acquisition, immunity and defence against infectious exogenous agents (hereafter called ‘parasites’ and including micro- and macroparasites) or proliferating indigenous organisms (Belkaid & Hand, 2014; Kamada *et al*., 2013; Round & Mazmanian, 2009) among others.

Nonetheless, the gut microbiota may differ greatly in natural environments between individuals, populations and species (Vujkovic-Cvijin *et al*., 2020). Its composition is even subject to high temporal variation for a given individual.

These variations are shaped by ecological and/or evolutionary processes, among which stochasticity, migration and/or adaptive differences in microbes (McDonald *et al*., 2020; Kolodny & Schulenburg, 2020). They are mediated by environmental features (e.g. acquisition of microorganisms from the environment, potentially through diet Ley *et al*., 2008; Moran *et al*., 2019), host factors (notably phylogeny, genetics or vertical transmission from mother to offspring) and interactions between hosts and their environment across space and time. For example, disruption of host-associated gut microbiota (termed “dysbiosis”) may occur as a result of environmental change and stress affecting the host. This has been shown in the context of anthropogenic pressures (e.g. chemical exposures, Rosenfeld, 2017; urbanisation, Stothart *et al*., 2019) or parasite infections (Trevelline *et al*., 2019).

Understanding the relationships between gut microbiota and parasites is crucial regarding their potential impacts on human and animal health (Clemente *et al*., 2012). Among the numerous studies of vertebrate microbiota, some of them have put an emphasis on the gut bacterial microbiota (called hereafter ‘gut bacteriota’) and their interactions with gastro-intestinal helminth parasites. On one hand, the gut bacteriota may act as an innate immune barrier to intestinal infections and influence the local colonisation and growth of eukaryotic parasites, including helminths, through competitive metabolic interactions or induction of host immune responses (Leung *et al*., 2018). On the other hand, helminth infections may also directly or indirectly affect the composition of the gut bacteriota via physical contact, competition for resources or host immunoregulation (see Kreisinger *et al*., 2015).

Interactions between helminths and the gut bacteriota may be positive or negative (Loke & Lim, 2015). They may lead to potentially local but also systemic physiological changes affecting host health. For example, helminth infections can lead to malnutrition and weight loss through the dysfunction of microbial metabolism that could result from negative impacts on fermentative gut bacteria (Leung *et al*., 2018). Besides, some helminth infections promote higher abundance of gut bacteria that produce short-chain fatty acids from dietary fiber (Zaiss *et al*., 2015). These metabolites circulate throughout the body and are important regulators of host physiology (glucose and fat metabolism) and immune system (Honda & Littman, 2016; Kim, 2021). Interactions between these helminths and gut bacteria may here increase the host anti-inflammatory and regulatory T cell suppressor responses, what may in turn affect host susceptibility to other infections as well as the outcomes of infections (Glendinning *et al*., 2014).

The gut microbiota may also influence microparasite infections through their immune function against local pathogenic bacteria colonization and their role in maintaining the intestinal epithelium integrity (Khosravi & Mazmanian, 2013). There is also strong evidence for systemic interactions between the gut microbiota and extra-intestinal microbiota communities, at least in laboratory mice (e.g. Rosshart *et al*., 2017;). This systemic impact of gut microbiota is mediated by host immunity (Zheng et al., 2020). As such, the gut microbiota produces metabolites (e.g., bacteriocins, short-chain fatty acids, microbial amino-acids) that translocate from the intestinal lumen to various organs (e.g., liver, brain, lung) through the circulatory system. This may induce tissue-specific immune responses, and affect the host’s susceptibility/resistance to (non enteric) pathogens (Winckler & Thackray, 2019; Pfeiffer & Sonnenburg, 2011). Most of these studies have focused on viruses (e.g., influenza A, coronaviruses, Karst & Wobus, 2019) and not yet on pathogenic bacteria (but see Rolhion & Chassaing, 2015). The systemic impact of gut bacteriota on microparasite infections still represents a fundamental knowledge gap in wild animals (Pascoe *et al*., 2017).

The three-way interactions between host’s gut bacteriota, gastro-intestinal helminths and microparasites have been scarcely investigated in a single system, despite clearly becoming pivotal in disease ecology. Yet, the growing interest on gut bacteriota/parasitism relationships in recent literature (P. T. Johnson *et al*., 2015) highlights the critical need for further empirical works. One main reason is the relatively low concordance of findings between previous studies – even for comparable host-parasite systems (e.g. for *Trichuris* sp and the gut microbiota, see Cortes et al., 2019; Lawson et al., 2021). Up to now, most of the research on this topic have been conducted on model species in laboratory settings. Although experiments under controlled conditions may help deciphering the mechanisms underlying these interactions between gut bacteriota and parasites in vertebrates (Pascoe *et al*., 2017), they also have inherent limitations. On the one hand, they only included a restricted number of targeted parasites (usually helminths and/or microparasites). Consequently, they often omitted the potential effects of species interactions between and within parasite communities at the intra-host level (Telfer *et al*., 2010). Co-infections by helminths species have been noticed by parasitologists for decades (Montgomery & Montgomery, 1989; Haukisalmi & Henttonen, 1993). Yet, co-infections between highly divergent micro- and macroparasites are also recognized to be the rule in most hosts in natural environments. Indeed, wild animals may carry simultaneously a large number of bacteria, helminths, viruses (Hoarau *et al*., 2020). On the other hand, they are unable to include – and then capture the complexity of – the environmental conditions as drivers of the composition of gut bacteriota and of the exposure or sensibility to these latter (Adair & Douglas, 2017). From there, studies in natural contexts deserve strong consideration because these environmental factors may impact deeply the relationships between gut bacteriota and parasitism. Empirical studies should enable to highlight associations, what is a critical pre-requisite before defining hypotheses about potential interactions and their underlying mechanisms.

Here, we strived to bridge these gaps by assessing the variability of gut bacteriota diversity and composition in wild populations of the bank vole *Myodes glareolus*, which is a small mammal reservoir of a large number of infectious agents (e.g., Abbate *et al*., In_revision). We studied the relationships between its gut bacteriota, parasite infracommunities (focusing on gastro-intestinal helminths and pathogenic bacterial infections) and host and environmental factors that may either influence or indicate the health status of the host (e.g., proxies such as the body mass index (BMI)). The current study therefore addressed two main questions: (1) how is the structure (composition and diversity) of the gut bacteriota influenced by host and environmental factors? (2) does the structure of gut bacteriota also reflect associations with gastro-intestinal helminth and pathogenic bacterial communities?

## Material and methods

### Data collection

#### Bank vole sampling

Bank voles (*Myodes glareolus*) were trapped in summer, between late June and early September 2014 in forests located in four French localities (Table 1, Figure S1) distributed along a North-South transect in Eastern France. These localities are separated by 40 to 120 km from one another. The standardized trapping protocol used here was described in details in Dubois et al. (2018).

**Table 1.**
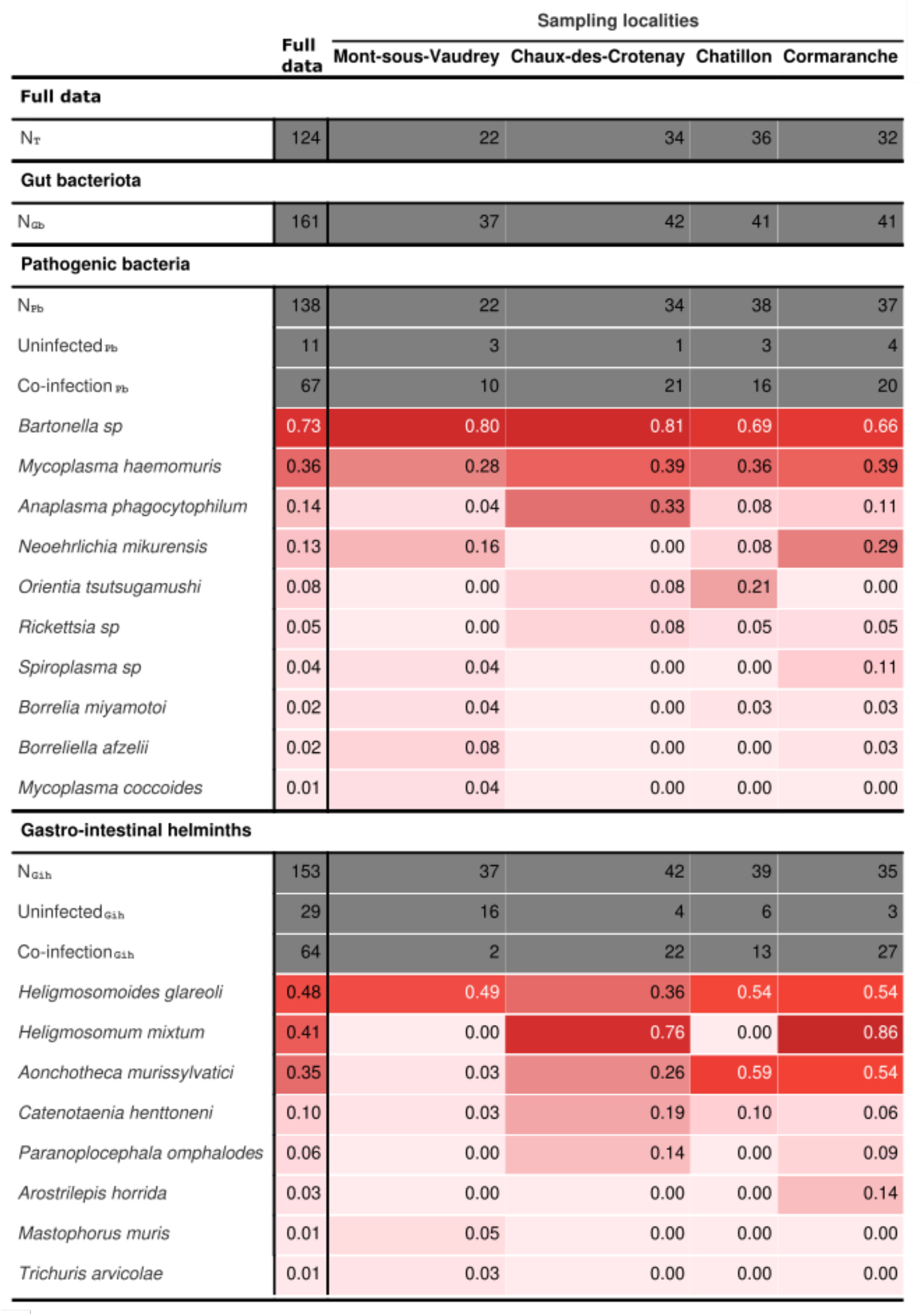
Number of bank voles analysed and prevalence of potentially pathogenic bacteria and gastro-intestinal helminths for each sampling locality. *N* is the number of bank voles analysed (Grey cells). *N*_*T*_ represents the number of individuals with data available for the three intra-host communities (gut bacteriota, pathogenic bacteria and gastro-intestinal helminths). *N*_*GB*_, *N*_*PB*_ and *N*_*GIH*_ respectively represent the number of individuals with data available for each of these intra-host communities. ‘*Uninfected’* corresponds to the number of uninfected bank voles for a given intra-host community. ‘*Co-infection’* corresponds to the number of bank voles infected with at least two parasites of a given intra-host community. Prevalence is provided for each pathogenic bacteria detected from the spleen, and each gastro-intestinal helminth. The red color gradient illustrates variations in prevalence (0% = light red to 100% = dark red).

Rodents were euthanized using isofluorane and cervical dislocation, as recommended by Mills (1995). A similar set of morphological measures was systematically recorded for each individual. Age groups (juveniles and adults) were defined according to body mass and sexual maturity. This latter was inferred using testes length and position, and seminal vesicle development for males, or uterus size for females. Body condition was estimated using the body mass index (BMI = weight/length^2^). The digestive tract and the spleen were removed and stored respectively in 96% ethanol and RNA later solution (−20°C).

Ethical statements: Animal capture and handling have been conducted according to the French and European regulations on care and protection of laboratory animals (French Law 2001-486 issued on June 6, 2001 and Directive 2010/63/EU issued on September 22, 2010). The CBGP laboratory has approval (D-34-169-003) from the Departmental Direction of Population Protection (DDPP, Hérault, France), for the sampling of rodents and the storage and use of their tissues.

#### Characterization of gut bacteriota

We first characterized the gut bacteriota of bank voles. We focused on the colon as rodent gut microbiota exhibits the highest level of bacterial diversity in the lower segments of the digestive tract (Suzuki & Nachman, 2016). DNA was extracted in 2016 from a 5 mm piece of colon tissue (taken about 1 cm far from the caecum - lumen was removed) of each bank vole using the ZymoBiomics 96 DNA Kit (Zymo) following the manufacturer’s instructions. We amplified a 251-bp portion of the V4 region of the 16S rRNA gene (16S-V4F [GTGCCAGCMGCCGCGGTAA] and 16S-V4R [GGACTACHVGGGTWTCTAATCC]), following Kozich et al. (2013) and as described in Galan et al. (2016). Samples were multiplexed using dual-indexes (index i5 in the forward primer and index i7 in the reverse primer). Negative controls for extraction (whole reagents without DNA), for PCR (PCR mix without DNA), and for indexing (wells without reagents corresponding to particular dual-indexes combinations). All DNA extractions were analysed twice using two independent technical replicates of amplicon libraries. PCR products were pooled, migrated and excised on a low melting agarose gel (1.25%) then purified using the NucleoSpin Gel and PCR Clean-Up kit (Macherey-Nagel) and quantified using the KAPA library quantification kit (KAPA Biosystems) standardized to 4nM by qPCR spectrophotometry (assay). Sequencing was performed on a 251-bp paired-end Illumina MiSeq run. The raw sequence reads (.fastq format) have been deposited in the Zenodo Repository.

Sequence data were processed as described in Galan et al. (2016) using the pipelines implemented in FROGS (Find Rapidly OTU with Galaxy Solution, Escudié *et al*., 2018). Briefly, the paired-end sequences were trimmed with CUTADAPT (Martin, 2011), merged with FLASH (MAGOC & SALZBERG, 2011), and clustered into fine-scale molecular operational taxonomy OTU units at 97% identity using the SWARM algorithm (Mahe *et al*., 2014) executed with aggregation parameter distance *d*=1 and a second pass performed on the seeds of previous clusters with *d*=3. As such, OTUs do not correspond to a fixed clustering threshold. Putative chimeras were removed using VSEARCH tools with de novo VUCHIME and the cross-validation method. Taxonomy was assigned with BLASTN+ (Camacho *et al*., 2009) using the SILVA SSU Ref NR 128 database as a reference (http://www.arb175silva.de/projects/ssu-ref-nr/). Filtering for false positives was carried out as proposed by Galan et al. (2016). In short, we discarded positive results associated with sequence counts below two OTU-specific thresholds, which checked respectively for cross-contamination between samples (using the negative controls for extraction and PCR) and incorrect assignment due to the generation of mixed clusters on the flow cell during Illumina sequencing, using a false index-pairing rate for each PCR product of 0.02%, based on estimates from Galan et al. (2016). For each sample, only OTUs found in the two technical replicates were considered as positive, and OTUs found in only one of the two replicates were removed. The number of sequences obtained for each technical replicate from a sample were summed.

Lastly, we discarded OTUs and samples containing less than 500 reads in the dataset, as well as OTUs considered to be contaminants, following (Salter *et al*., 2014). Number of reads per OTU were finally normalised to proportional abundance within each rodent (McKnight *et al*., 2019). We only considered the family taxonomic rank for further analyses, but analyses at the phylum level provided similar results (not shown).

#### Detection of pathogenic bacteria and gastro-intestinal helminths

We described the presence/absence of pathogenic bacteria from the spleen of each bank vole. This lymphoid organ filters microbial cells in mammals and as such, enables to recover recent infections (Abbate *et al*., In_revision; Diagne *et al*., 2017). Molecular protocols, bioinformatics pipelines and data filtering were similar to those described above (gut bacteriota), except for the DNA extraction from splenic tissue using DNeasy 96 Tissue Kit (Qiagen). The potential pathogenicity of each bacterial OTU was assessed based on published literature and on the Gideon database (https://www.gideononline.com/). Opportunistic pathogens (*i*.*e*. commensal agents in healthy hosts, that become pathogenic when the balance of the immune system is disrupted) were discarded from the dataset. Only the information of the presence / absence of pathogenic OTUs was considered. For each bank vole, helminths were carefully extracted and counted from the different sections of the digestive tract (stomach, small intestine, large intestine and caecum), and classified by morphotype then stored in 95% ethanol for further accurate identification. The latter was based on unambiguous morphological criteria using conventional microscopy and generalist identification keys or specific literature when available (Anderson *et al*., 2009 ; Khalil *et al*., 1994 ; Ribas Salvador *et al*., 2011).

#### Statistical analyses

All statistical analyses were implemented in R v4.0.3 (team, 2020). For more convenience, gut bacteriota, pathogenic bacteria and gastro-intestinal helminths were further described as ‘intra-host communities’.

#### Gut microbiota diversity and composition

Description and analyses of bacterial communities were performed using the PHYLOSEQ package (McMurdie & Holmes, 2013). We considered three features to analyse within hosts’ gut microbiota. i) We looked for enterotypes, *i*.*e*. distinct community composition types of gut bacteriota, as found in humans (Arumugam *et al*., 2011; Holmes *et al*., 2012), using the Dirichlet Multinomial Mixtures DMM (Morgan, 2021). ii) We analysed the Firmicutes /Bacteroidetes (F/B) log-ratio, as it is often used as a proxy of health or metabolism in humans and mice (Ley *et al*., 2005; Toumi *et al*., 2022). We calculated this ratio with the MICROBIOTA package (Lahti & Shetty, 2017). iii) We characterized the alpha diversity using two metrics, the specific richness (*i*.*e*. number of taxa within the host individual) and the Shannon index as recommended in (Haegeman *et al*., 2013).

We estimated the beta diversity, *i*.*e*., the dissimilarity between host individuals in their gut bacteriota using Bray-Curtis distances. We considered the relative abundance of OTUs (family).

Influence of host and environmental factors on gut bacteriota diversity and composition We tested the influence of individual characteristics (age class, gender, BMI) and localities, independently on the F/B log-ratio and on the alpha diversity using generalized linear models (GLM). We considered a negative binomial error distribution for the F/B ratio and the specific richness, and a gaussian distribution for the Shannon index. Best model selection was performed considering models with all possible combinations of factors and the DREDGE function of the MUMIN package. The best model was selected using the Akaike information criterion corrected for small sample size AICc, (J. B. Johnson & Omland, 2004). We assessed the effect of each factor in the best model with the ΔAICc index. When the factor locality was significant, Tukey’s post-hoc tests were applied to evaluate pairwise differences between localities, using the MULTCOMP package (Hothorn *et al*., 2008). Residuals were checked to graphically to ensure that all assumptions regarding normality, independence and the homogeneity of variance were satisfied.

We evaluated the influence of geographic distance on the dissimilarities in gut bacteriota by performing Mantel tests and using Pearson correlation (10,000 permutations). These tests have less statistical power to address questions related to the variation in community composition data among sites. Therefore, we also analysed the factors shaping the dissimilarities in gut microbiota composition using several functions of the VEGAN package (Oksanen *et al*., 2020). Distance-based redundancy analyses (db-RDA) were performed to analyse the effect of individual explanatory factors (age class, gender, BMI) and sampling localities on dissimilarities in gut microbiota composition. Redundancy analyses are appropriate to test hypotheses about the origin and maintenance of the variation in β diversity (Legendre et al. 2005). We used the CAPSCALE function, followed by permutational multivariate analyses of variance (PERMANOVA). We selected the best model, *i*.*e*., the most parsimonious one, using the ORDIR2STEP function (*P*-value adjusted and *R*^*2*^ adjusted). For each factor, we evaluated the intra-group dispersion using the BETADISPER function as PERMANOVA analyses are sensitive to differences in dispersion among groups. A Tukey’s test was done to see if and which groups differed in relation to their variances. Lastly, we used DESEQ2 package (Love et al., 2014) to identify the changes in bacteria taxa that best explained gut bacteriome dissimilarities between individuals and localities. We performed GLMs with negative binomial error (NBINOMWALDTEST method) and significant differences were obtained after Benjamini & Hochberg corrections. They were visualised using the METACODER package (Foster et al., 2017).

#### Relationships between gut bacteriota and pathogenic communities

We estimated the alpha diversity of the gastro-intestinal helminhth and pathogenic bacteria community using the richness index (presence/absence data). We used GLMS and model selection process described above to analyse whether the alpha diversity of each intra-host community (gut bacteriota and pathogenic communities) was influenced by the alpha diversity of the two other ones.

We estimated the beta diversity of the gastro-intestinal helminth and pathogenic bacteria community using the Jaccard index (presence/absence data). The relationships between intra-host community dissimilarities were investigated using three approaches. i) We applied partial Mantel tests using MULTI.MANTEL (phytools package Revell, 2012) to analyse the correlation between two matrices of dissimilarities (corresponding to two different communities), while controlling for the effect of a third dissimilarity matrix (third community). ii) We used db-RDA to analyse more deeply the relationships between the gut bacteriota and the pathogenic (bacteria and helminths) communities. We included the alpha diversity indices (richness specific) and infectious status as presence / absence) of pathogens with prevalence greater than 10% in at least one locality as explanatory variables in these analyses. We selected the best model using the ORDIR2STEP method. iii) We used DESEQ2 to determine the gut bacteria taxa whose relative abundances changed with significant explanatory variables.

## Results

A total of 186 bank voles were trapped during the fieldwork campaign over the four targeted localities. For technical reasons (e.g., poor sample preservation, missing data), we could study the three intra-host communities for 124 rodents only.

### Characterization of the gut bacteriota: taxa and enterotypes

Once the quality control steps were applied, the gut bacteriota dataset included 161 bank voles. We detected 10 phyla and 61 families of bacteria. At the phylum level, we found six predominant taxa that represented 99% of the gut bacteria relative abundance (Figure S2). At the family level, 11 families represented 93% of the relative abundance of the gut bacteriota (Figure 1A).

**Figure 1.**
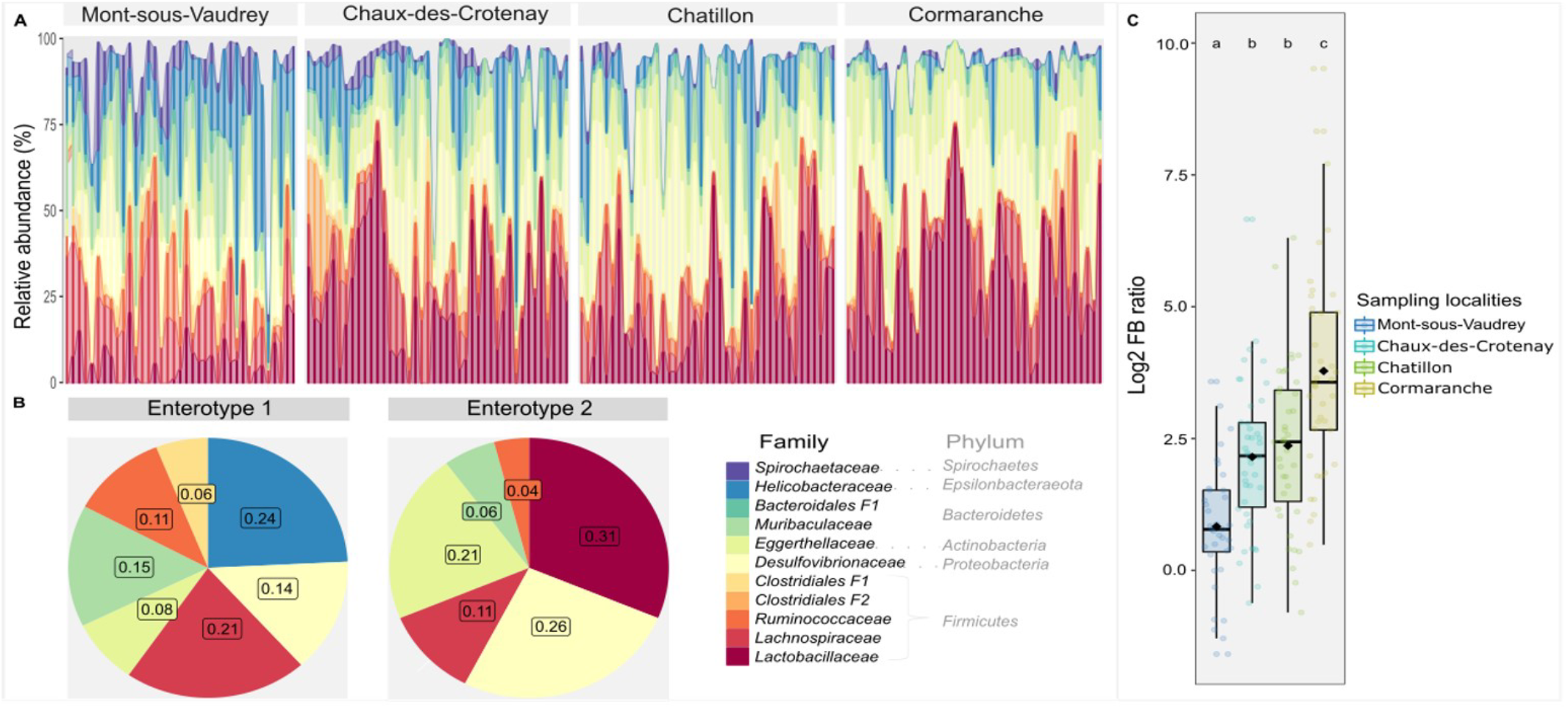
Composition of the intestinal bacteriota. A) The bar plot shows the individual variations of 11 bacterial families (F= Unknown family) belonging to 6 phyla and representing 93% of the total composition. Individuals (bars) are grouped by sampling localities, which are ordered from North to South. Each color represents a taxa. B) The composition of the two enterotypes identified using Dirichlet multinomial mixtures (DMMs), at family rank, is shown. Bacterial families are represented using the same colors as in A. C) the ratio (Firmicutes / Bacteroidetes) is shown for each sampling locality. Box and whisker plots represent the median and interquartile values. Black dots correspond to the mean value, and colored dots correspond to individuals.

Different letters indicate statistically significant differences at P < 0.05, with pairwise Tukey post hoc adjustments.

We distinguished two enterotypes from the DMM approach. One (enterotype 1) was mainly composed of the families *Helicobacteraceae, Lachnospiraceae, Muribaculaceae and Desulfovibrionaceae*, while the other (enterotype 2) mainly included *Lactobacillaceae, Desulfovibrionaceae* and *Eggerthellaceae* (Figure 1B).

### Diversity of the gut bacteriota : the influence of sampling locality and host condition

We found that the *Firmicutes / Bateroidetes* ratio varied significantly between localities (Figure 1C). Overall, northern localities exhibited lower F/B ratio than southern ones, with all pairwise comparisons being significant except Chatillon *versus* Chaux-des-Crotenay. Individual characteristics did not influence this ratio (Table S1).

The sampling locality had a significant global effect on the alpha diversity of the gut bacteriota (*GLMs*. Specific richness: *F* = 8.49, *P* < 10^−3^; Figure 2A; Shannon index: *F* = 4.74, *P* = 3 × 10^−3^; Figure 2B; Table S2A). The locality Cormaranche exhibited a higher specific richness than all other localities (*Tukey post hoc test*. Mont-sous-Vaudrey: Z = 5.13, *P*_adj_ < 10^−3^, Chaux-des-Crotenay: *Z =* 4.57, *P*_*adj*_ < 10^−3^ and Chatillon: *Z* = 3.62, *P*_*adj*_ = 1.7 × 10^−3^) but a lower level of diversity than Mont-sous-Vaudrey when considering taxa relative abundance (*Tukey post hoc test*. Shannon index: *Z* = -3.64, *P*_adj_ = 1.5 × 10^−3^, Figure 2B). Body condition (BMI) was also found to have a significant effect, but only when considering specific richness (*t* = 2.91; *P* = 4 × 10^−3^) – with higher values of BMI associated with increasing species richness. All these results are detailed in Table S2A and Figures S3).

**Figure 2.**
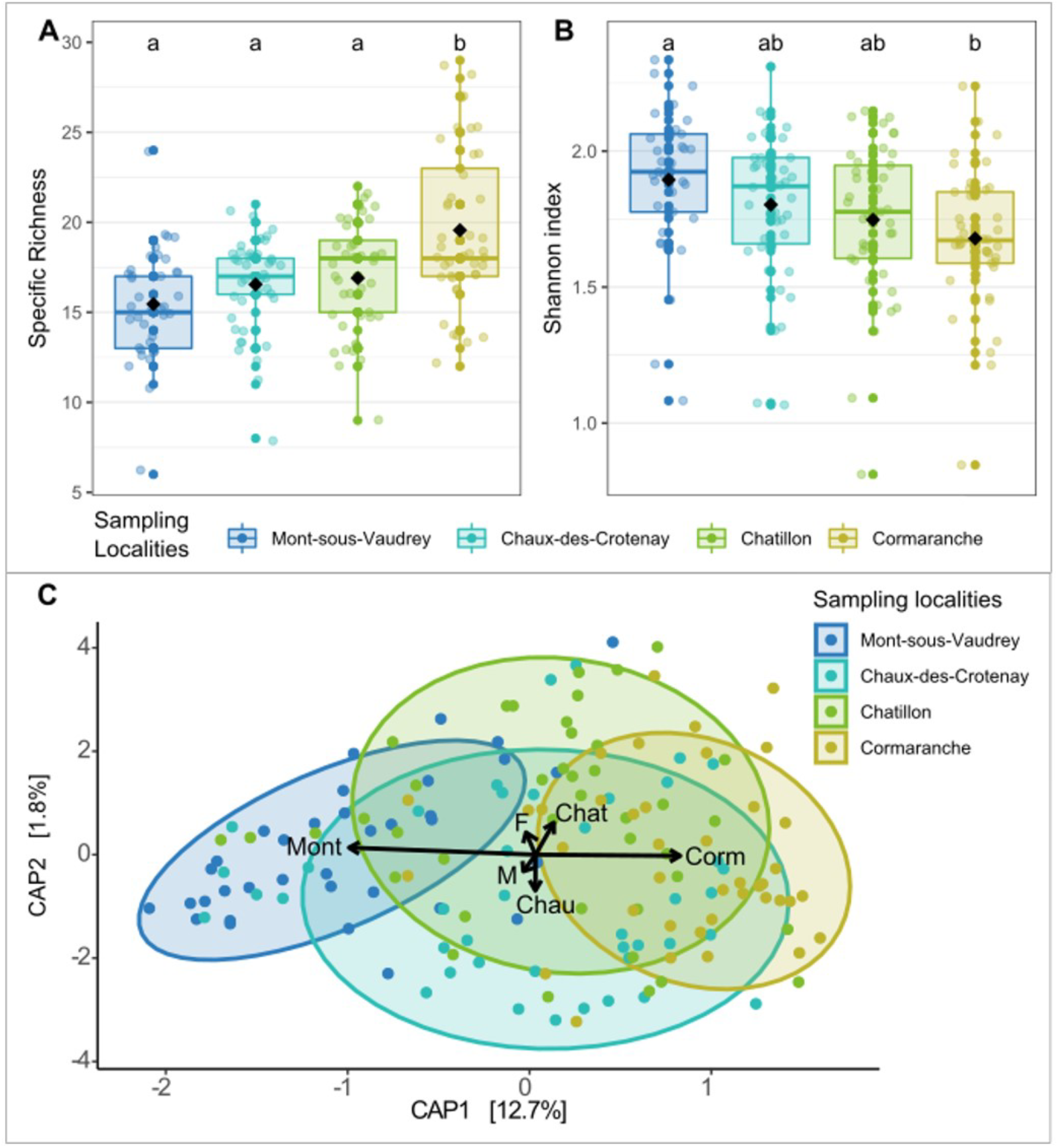
Variations of the gut bacteriota alpha diversity between localities. Alpha diversity is represented using A) the specific richness of the gut bacteriota, and B) the Shannon index of the gut bacteriota, Results are shown per locality, ordered from North to South. Each colored point represents an individual. Black points indicate the average alpha diversity per locality. Box-and-whisker plots represent the median and interquartile values. Different letters denote statistically significant differences at *P* < 0.05, with pairwise post-hoc Tukey adjustments. C) Distance-based redundancy analysis (db-RDA) of the gut bacteriota (family) based on Bray– Curtis dissimilarities. Dots represent individuals. The colors and shapes of the dots are associated with different factors : Localities from North to South (Mont-sous-Vaudrey: Mont, Chaux-des-Crotenay: Chau, Chatillon: Chat and Cormaranche: Corm) and gender (females: F and males: M). Significative factors based on the ordiR2step analysis are shown as arrows. Ellipses represent a 80% confidence interval around the centroid of the clusters, for each locality.

### Composition of the gut bacteriota: sampling locality as the main factor of variation

We found a significant positive relationship between the dissimilarities in gut bacteriota composition and the geographic distance (*Mantel test. r* = 0.25, *P* = 10^−4^, Table S3A).

We found a significant effect of the sampling localities (*db-RDA. R*^*2*^_*adj*_ = 0.16, *P* = 1 × 10^−3^) and host gender (*db-RDA. R*^*2*^_*adj*_ = 0.01, *P* =0.027) on gut microbiota composition. The CAP1 axis discriminated Mont-sous-Vaudrey and Cormaranche localities (12.7% of the total variance, Figure 2C). However, this result has to be taken cautiously as significant differences of data dispersion were detected between localities (*betadisper. P* = 1 × 10^−3^). The locality Cormaranche showed a lower dispersion compared to all other localities (Tukey multiple comparisons, Table S3B).

We detected significant differences in the relative abundance of specific taxa using DeSeq2 (Table 2; Table S3C). The main changes (Log2 fold values higher than 20) were detected between the northern (Mont-sous-Vaudrey) and southern localities. The northern population was involved in 75% of all significant pairwise differences (Log2 fold change in composition > 10). The gut bacteriota of these bank voles includes less Clostridiales (one unknown family; Firmicutes), Bifidobacteriaceae (Actinobacteria) and Desulfovibrionales (two unknown families; Proteobacteria), but more Erysipelotrichaceae (Firmicutes) than in all the three other southern populations. The gut bacteriota of bank voles from Cormaranche (South) is characterized by less Erysipelotrichaceae (Firmicutes), than in the three northern populations.

**Table 2.**
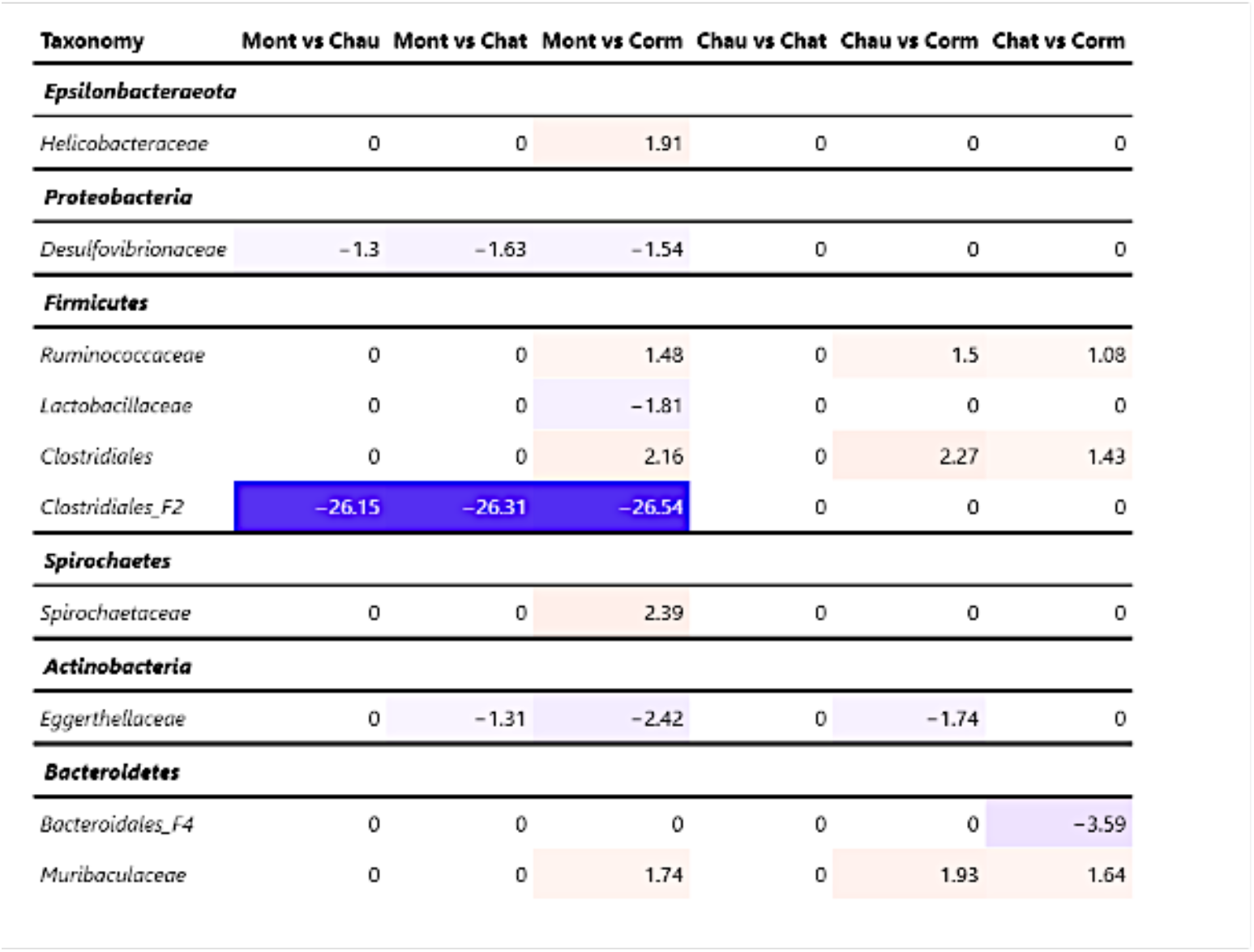
Pairwise comparisons of the relative abundance of the gut bacteriota between sampling localities. Mont = Mont-sous-Vaudrey, Chau = Chaux-des-Croteay, Chat = Chatillon, Corm = Cormaranche. The Log^2^ fold value is indicated for significant changes in abundance between two localities. Blue and red colors respectively correspond to negative and positive values. Higher absolute changes in Log^2^ fold are emphasized with darker colors. The notation “order_fx” or “class_fx” is used when there was no assignation at the family level with the SILVA database. Phylum is indicated in bold.

Differences in the composition of the gut bacteriota between males and females bank voles were driven by the phylum *Firmicutes*, with males exhibiting higher relative abundance of this taxa than females (Table S3C).

### Relationships between the diversity of the three intra-host communities

We found a significant relationship between the specific richness of the gut bacteriota and the richness of the helminth community. A more diverse gut bacteriota was associated with a greater number of helminth species infecting bank voles (*GLM. F =* 14.09, *P* < 10^−3^; Figure 3A; Table S2B). We also found a positive relationship between the specific richness of the pathogenic bacteria and the richness of the gastro-intestinal helminth community (*GLM. F*= 6.99, *P* = 9 × 10^−3^; Figure 3A; Table S2B). Lastly, we found a significant effect of the specific richness of both the gut bacteriota and of pathogenic bacteria on the richness of the gastro-intestinal helminth community (*GLM*. Gut bacteriota. *t* = 3.50, *P* < 10^−3^; pathogenic bacteria *t* = 2.38, *P* = 0.019; Figure 3A, Table S2B).

**Figure 3.**
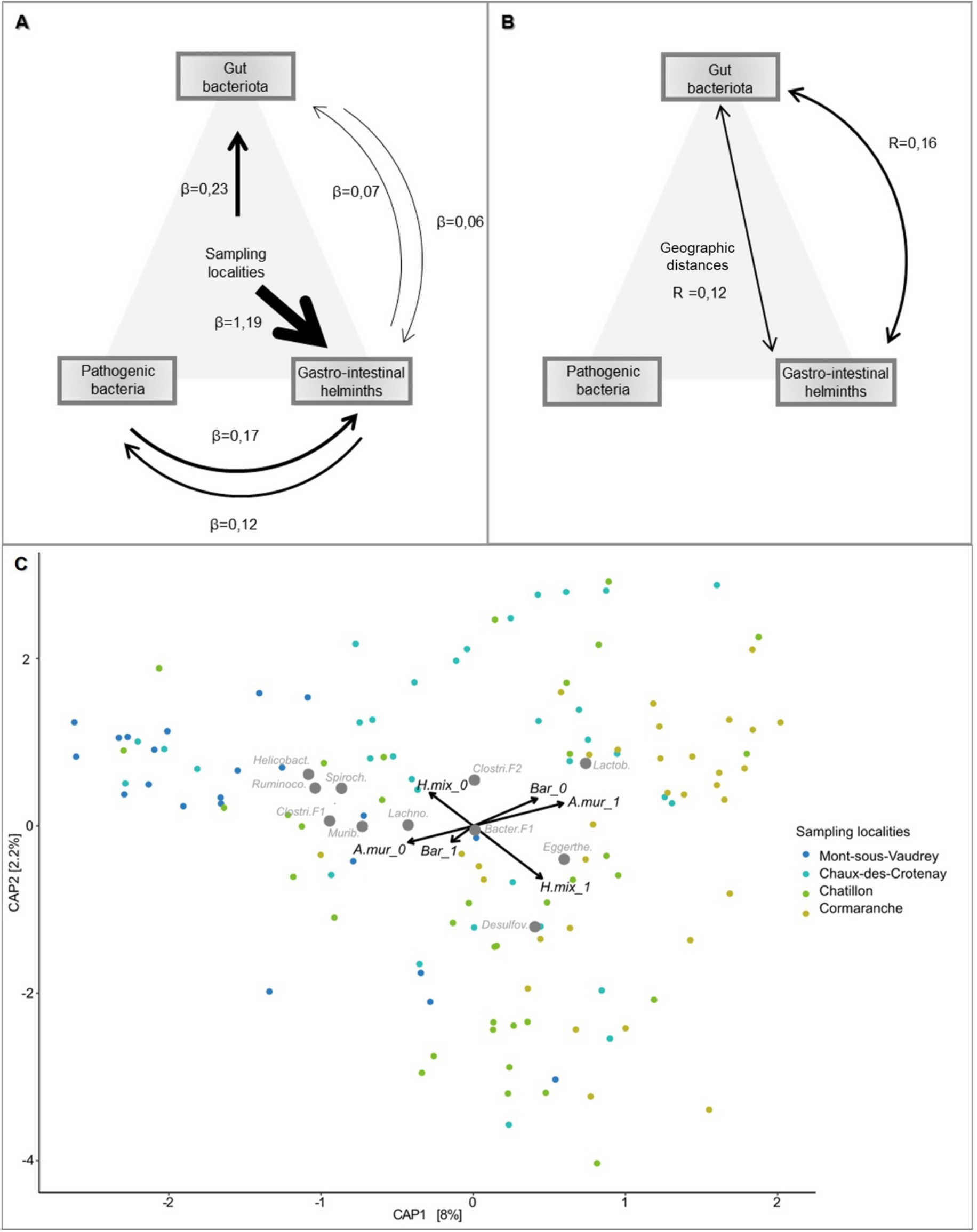
Associations between the diversity and composition of the three intra-host communities. The two upper diagrams show the relationships between A) the specific richness and B) the composition of the three communities, while taking into account the influence of the sampling localities or distance geographic. The effect size and the direction of the relationship between communities are represented using an arrow, its width corresponding to the estimate (β) x 10 or the correlation index *R* x 10. Only significant effects are represented. C) This db-RDA triplot shows the structure of the gut bacteriota and the correlations with the pathogen communities. The arrows correspond to the significant explanatory variables. Each point corresponds to an individual, and the colors correspond to the different sampling localities. A.mur: *Aonchotheca mzrissylvatici*, Bar: *Bartonella* sp, H.mix: *Heligmosomum mixtum*.; Helicobacter.: Helicobacteraceae, Spiroch.: Spirochaetaceae, Clostri.F2: Clostridiales_f2, Clostri.F1: Clostridiales_f1, Lactob.:Lactobacillaceae,Eggerthe.: Eggerthellaceae, Desulfov.: Desulfovibrionaceae, Bacter.F1: Bacteroidales_f1, Lachno.:Lachnospiraceae; Murib.: Muribaculaceae, Ruminoco.: Ruminococcaceae

### Relationships between the composition of the three intra-host communities

We found a positive relationship between dissimilarities in the gut bacteriota and dissimilarities in the gastro-intestinal helminth community composition (*partial Mantel test. R* = 0.16, *P* =1 × 10^−4^; Figure 3B), but not with dissimilarities in the pathogenic bacteria community composition (*partial Mantel test. r* = 0.02, *P* = 0.28). After controlling for geographic distances, dissimilarities in gut bacteriota composition remained significantly correlated with dissimilarities in helminth community composition (*partial Mantel test. R* =0.12; *P* = 0.001). Further details are provided in Table S3A.

We detected significant associations between the whole composition of the gut bacteriota and the presence / absence of three pathogens: *Aonchotheca murissylvatici* (*ordistep db-RDA. R*^*2*^_*adj*_ = 0.0, *P* = 2 × 10^−3^), *Bartonella* sp (*ordistep db-RDA. R*^*2*^_*adj*_ = 0.01, *P* = 0.04) and *Heligmosomum mixtum* (*ordistep db-RDA. R*^*2*^_*adj*_ = 0.06, *P* = 2 × 10^−3^). The db-RDA triplot based on the two first axes only represented 10.2% of the total variance (Figure 3C; Figure S4). It showed that individuals infected with *Aonchotheca murissylvatici* or *Heligmosomum mixtum*, but not infected with *Bartonella sp*., had more *Lactobacillaceae* (*Firmicutes*), *Desulfovibrionaceae* (*Proteobacteria*) and *Eggerthellaceae* (*Actinobacteria)*. These individuals also had less *Spirochaetaceae* (*Spirochaeta), Muribaculaceae (Bacteroidetes), Helicobacteraceae* (*Epsilonbacteraeota*) and *Ruminococcaceae* (*Firmicutes*). This pattern is correlated with the sampling localities. Individuals from northern localities are distributed on the left side of the CAP1 axis, and southern ones on the right side of it (Figure 3C). Neither the specific richness of pathogenic bacteria nor the specific richness of the gastro-intestinal helminth community had a significant effect on the global composition of the gut bacteriota (Table S3D).

The specific richness of the gastro-intestinal helminth community, as well as infections with *A. murissylvatici* and *H. mixtum*, were only slightly associated with different relative abundance of particular gut bacteria taxa (*DeSeq2*. Log2 fold changes did not exceed 3.5). These changes concerned four main families. Rhizobiaceae and Spirochaetaceae showed negative associations with gastro-helminth specific richness and *A. murissylvatici*. Mollicutes (undetermined family) and Saccharimonadaceae showed positive associations with *A. murissylvatici* and *H. mixtum* (Table 3). More details are provided in Table S3E.

**Table 3.** Changes in relative abundance of the gut bacteriota with regard to infectious status (helminths and pathogenic bacteria) and specific richness. The Log^2^ fold change in relative abundance is indicated for significant values only. Negative values are represented with blue colors, positive values with red colors. Higher absolute changes in Log^2^ fold are emphasized with darker colors.

| Phylum | Family | log2FoldChange |
| --- | --- | --- |
| **Richness of GI helminths** |  |  |
| Actinobacteria | Eggerthellaceae | 0.67 |
| Proteobacteria | Rhizobiaceae | -3.79 |
| Epsilonbacteraeota | Helicobacteraceae | -0.89 |
| **Aonchotheca murissylvatici** |  |  |
| Proteobacteria | Desulfovibrionaceae | 1.13 |
| Tenericutes | Mollicutes_f1 | 3.07 |
| Patescibacteria | Saccharimonadaceae | 3.03 |
| Spirochaetes | Spirochaetaceae | -1.75 |
| Actinobacteria | Eggerthellaceae | 1.4 |
| **Heligmosomum mixtum** |  |  |
| Tenericutes | Mollicutes1 | 3.34 |
| **Mastophorus muris** |  |  |
| Proteobacteria | Desulfovibrionales3 | -29.98 |
| Firmicutes | Clostridiales_f1 | -21.41 |
| Firmicutes | Clostridiales_f2 | -23.73 |
| **Catenotaenia henttoneni** |  |  |
| Proteobacteria | Rhizobiaceae | -24.71 |
| Proteobacteria | Burkholderiaceae | -23.96 |
| Firmicutes | Erysipelotrichaceae | -20.6 |
| **Paranoplocephala omphalodes** |  |  |
| Firmicutes | Clostridiales_f2 | -25.37 |
| Proteobacteria | Desulfovibrionales_f3 | -23.74 |
| Proteobacteria | Rhizobiaceae | -24.86 |
| Proteobacteria | Burkholderiaceae | -22.69 |
| Firmicutes | Erysipelotrichaceae | -22.4 |
| Firmicutes | Christensenellaceae | -5.24 |
| **Trichuris arvicolae** |  |  |
| Firmicutes | Christensenellaceae | -20.69 |
| Firmicutes | Clostridiales_f1 | -18.62 |
| Bacteroidetes | Bacteroidales_f4 | -18.38 |
| **Richness of Pathogenic bacteria** |  |  |
| Bacteroidetes | Bacteroidales_f3 | -28.51 |
| **Neoehrlichia mikurensis** |  |  |
| Firmicutes | Erysipelotrichaceae | -23.42 |
| Firmicutes | Clostridiales | -2.72 |
| Bacteroidetes | Muribaculaceae | -1.76 |
| **Orientia tsutsugamushi** |  |  |
| Firmicutes | Erysipelotrichaceae | -22.95 |
| Proteobacteria | Desulfomicrobiaceae | -5.71 |
| Bacteroidetes | Muribaculaceae | -2.45 |
| **Anaplasma phagocytophilum** |  |  |
| Firmicutes | Ruminococcaceae | -1.75 |
| **Rickettsia sp** |  |  |
| Firmicutes | Erysipelotrichaceae | -21.81 |

In the opposite, we found strong changes in relative abundance of gut bacteriota families with other gastro-intestinal helminths and pathogenic bacteria infections (*DeSeq2*. Log2 fold higher than 18, Table 3). It concerned infections with the helminth species *Mastophorus muris, Catenotaenia henttoneni, Paranoplocephala omphalodes* and *Trichuris arvicolae*.

These associations were all negative and mostly involved the same bacterial families (Table S3E), namely undetermined families of Bacteroidales (Bacteroidetes), Desulfovibrionales (Proteobacteria) or Clostridiales (Firmicutes), Erysipelotrichaceae (Firmicutes), Rhizobiaceae (Proteobacteria) and Burkholderiaceae (Proteobacteria).

Considering pathogenic bacteria, we found that higher levels of specific richness were associated with lower relative abundance of an undetermined Bacteroidales family (Bacteroidetes), and that *Neoehrlichia mikurensis, Orientia tsutsugamushi* and *Rickettsia* sp infections were associated with strong decreases in relative abundance of *Erysipelotrichaceae* (Firmicutes) (Table 3; Table S3E). Other associations between bacterial infections and changes in relative abundance of specific gut bacteriota taxa were detected, but with little size effect (*DeSeq2*. log2 fold changes lower than 5).

## Discussion

Understanding the complex interlinkages between host microbiota, host-pathogen interactions and health in wild animal populations has become a key topic in disease ecology. Such understanding is instrumental for deciphering population dynamics, and designing strategies for zoonotic risk management or biodiversity conservation. Here, we use a combination of metabarcoding and community ecology approaches to *(i)* describe the gut microbiota of wild rodent populations and their variations at a regional geographical scale, and *(ii)* explore the three-way relationships between the gut bacteriota and communities of gastro-intestinal helminths and pathogenic bacteria.

### Spatial variations of gut bacteriota and their potential causes

The gut microbiota of bank voles has been mainly examined in the context of exposure to radioactive pollutants (e.g., Lavrinienko (2018), but see Knowles et al. (2019)). In this study, we focused on localities sampled at a regional scale (100 km) along a North-South gradient in Eastern France.

We found significant inter-individual variations in the gut bacteriota composition although host factors such as gender and age played little role. Interestingly, we found that all individuals were clustered within two groups, which could be distinct enterotypes if longitudinal surveys confirmed this clustering through time (Arumugam *et al*., 2011).

Enterotypes have already been described in wild rodents (Goertz *et al*., 2019; Li *et al*., 2016), and they might reflect distinct ways of generating energy from substrates available in the digestive tract, as well as differences in diet (Rinninella *et al*., 2019; Wang *et al*., 2014).

These enterotypes could be associated with different diets in bank voles: one oriented toward seeds and plants and another one toward insects and berries. Indeed, enterotype 1 is characterized by families (namely, Helicobacteraceae, Lachnospiraceae and Muribaculaceae) that are involved in the breakdown of carbohydrates, fermentation of plant saccharid and degradation of glycan (see refs in Goertz *et al*., 2019). These families are also predictive signals of a high-fat diet in mice (Bowerman *et al*., 2021; Rodriguez-Daza *et al*., 2020).

Conversely, enterotype 2 is characterized by families (namely, Lactobacillae and Eggerthellaceae) that can be involved in the digestion of fermented food (e.g., rodent food store over winter), and insect skeleton (see refs in Maurice *et al*., 2015) or in the degradation of polyphenol (Rodriguez-Daza *et al*., 2020). All these aliments have varying nutritional and chemical composition and may be part of bank vole diet (Ecke *et al*., 2018). The fraction of these different types of resources in bank vole diet may vary with resource preference or availabilities, reproductive status, sampling date and location (e.g., Maurice *et al*., 2015). It would be interesting to develop semi-natural experiments to survey rodent diet and gut microbiome through time, and analyse the link with enterotypes in bank voles (Wang *et al*., 2014).

There are now many evidence that the environment in which hosts evolve (through abiotic and biotic factors) is likely to shape variations in gut bacteriota composition between localities sampled and studies. Previous works have already shown that the structure of rodent gut microbiota varied between localities at large spatial scales due to biogeographic or genetic factors (Linnenbrink *et al*., 2013). Geographic variability has also been found at smaller spatial scales (e.g., few km Goertz *et al*., 2019). Here, our results provide significant evidence for spatial structure of gut bacteriota between bank vole populations that are between 50 and 130 km away, with no clear barrier to dispersal or gene flow (Dubois *et al*., 2018).

We observed gradual changes between bank voles from the northern and southern populations in terms of gut bacteriota richness, evenness, composition and particularly Firmicutes/Bacteroidetes ratio. Although the links between the diversity and functional capacity of the gut bacteriota still need to be better understood (Worsley *et al*., 2021), it is largely assumed that changes in diversity are associated with shifts in metabolism (Reese & Dunn, 2018). Bank voles from southern populations exhibit higher specific richness and lower evenness of the gut bacteriota, as well as lower dispersion of gut bacteriota composition. They have higher levels of body condition and F/B ratio, which are indicative of an optimisation of calorie intake and absorption, weight gain and fat storage (see refs in Wolf *et al*., 2021).

Altogether, these results could suggest strong constraints on gut bacteriota function to maximise energy extraction. The northern populations show the opposite patterns. Lower BMI and lower levels of F/B ratio might reflect energy production and conversion, amino acid transport and metabolism. Diversity patterns (higher evenness and lower specific richness of the gut bacteriota) could suggest lower stochasticity and/or directional selection. Further studies are required to investigate the eco-evolutionary processes driving these changes in gut bacteriota.

Lastly, these differences in gut microbiota composition between the northern and southern populations might also reflect physiological variations related to physiology, health and (potentially) immunity. Indeed, Clostridiales and Bifidobacteriaceae participate in the maintenance of intestinal homeostasis, and in the regulation of inflammation or in the gut barrier function (Arboleya *et al*., 2016; Hakansson & Molin, 2011; Lopetuso *et al*., 2013). Specific taxa within Erysipelotrichaceae may be correlated with inflammation or have immunogenic potential (Kaakoush, 2015; Zhai *et al*., 2019). Desulfovibrionales activities result in the production of H_2_S, which in turn negatively affects the gut barrier, production of endotoxins and pro-inflammatory cytokines (Hu *et al*., 2022). Our previous work revealed that bank voles from these southern populations had lower basal level of *Tnf-a* (a pro-inflammatory cytokine) and higher level of *Mx2* antiviral gene expression than those from these northern populations (Dubois *et al*., 2018). Future studies should assess *(i)* the potential relationships between variations in gut bacteriota composition and *(ii)* the capacities to regulate or mount immune responses and inflammation in these bank vole populations.

### Three-way relationships between intra-host communities

At the community level, we have not found strong evidence for three-way relationships between the gut microbiota (diversity or composition), the gastro-intestinal helminth and pathogenic bacteria communities. Neither the specific richness nor the global composition of a given community are related to the richness or composition of the two other ones. By contrast, particular taxa seem to be involved in these three-way relationships.

First, some infections are significantly associated with the global composition of the gut bacteriota, but have only little impact on specific gut bacterial taxa. This result concerns two helminths *Heligmosomum mixtum, Aonchotheca murissylvatici* and the hemotrophic bacteria *Bartonella*. Opposite patterns are observed for the helminths and for the bacteria. They could reflect the antagonistic impacts of these infections on the gut bacteriota, or negative interactions between these pathogens. On one hand, some evidence suggests that *Bartonella* may be acting as a symbiont more than a pathogen (McKee *et al*., 2021). Significant coevolutionary congruence has been found between *Bartonella* species and their rodent hosts, and *Bartonella* infections in rodents lead to an asymptomatic long lasting intra-erythrocytic bacteraemia (Deng *et al*., 2012; Lei & Olival, 2014). It would be interesting to test whether associations between *Bartonella* and gut bacteriota could corroborate the hypothesis of coadaptation between these bacteria and their rodent hosts (Hayman et al., 2013). On the other hand, some hookworms have been shown to induce changes in rodent gut bacteriota (review in Mutapi, 2015). Infection of mice with the nematode *Heligmosomoides polygyrus* (which is phylogenetically close from *Heligmosomum mixtum*) lead to an increased abundance of Lactobacillaceae in the gut microbiome (Reynolds *et al*., 2014), as suggested by the correlation detected in our study. Lastly, negative interactions between *H. mixtum or A. murissylvatici* and *Bartonella* are probable. Indeed gastro-intestinal hookworms are known to induce anaemia (Seguel & Gottdenker, 2017), while *Bartonella* invades and replicates in red blood cells. This resource limitation driven by helminths on erythrocyte-dependent infectious agents is an important driver of helminth-microparasite coinfection (Graham, 2008).

Therefore, the negative associations detected here between *Bartonella* and *H. mixtum or A. murissylvatici* – and their respective links with gut bacteriota composition – likely seem to be driven by potential complex antagonistic, synergistic and symbiotic interactions that need to be further explored.

Second, other infections are strongly associated with large changes in the relative abundance of one or few specific taxa from the gut bacteriota, but not with the global composition. Three bacterial infections (*Neoehrlichia* sp, *Orientia* sp and *Rickettsia* sp) as well as *P. omphalodes* infections exhibited the same pattern: they were associated with a lower relative abundance of Erysipelotrichaceae. It is striking to find such common associations for these infectious agents because observed changes in the gut microbiota during infection are rarely consistent, even for single pathogens (Sabey *et al*., 2021). The most obvious features shared between these infectious agents is that they are transmitted by arthropods. Unfortunately, knowledge on Erysipelotrichaceae and its links with infection or dysbiosis is still deficient to explain the pattern observed. To our knowledge, such associations have been investigated in humans only. They have shown that increased abundance of Erysipelotrichaceae could be associated with a number of diseases such as tuberculosis, HIV and norovirus infections, inflammation-related intestinal disease and metabolic disorders (Kaakoush, 2015). The reasons why *Neoehrlichia* sp, *Orientia* sp, *Rickettsia* sp or *P. omphalodes* infections are associated with a decreased abundance of Erysipelotrichaceae in bank voles remain to be investigated.

### Marked relationships between gut bacteriota and gastro-intestinal helminth communities

While we do not detect three-way relationships between the whole composition of gut bacteriota, helminth and pathogenic bacteria communities, we highlight strong pairwise associations between helminth community and gut bacteriota. The former pattern might be explained by the fact that the pathogenic bacteria detected here do not constitute a functional community. The main reason is that their ecological niche can be very different. For example, hemotrophic *Mycoplasma* parasitizes erythrocytes (Alabi *et al*., 2020) while *Borrelia* disseminates through the bloodstream and/or lymphatic system to invade and colonize various tissue (Zeidner *et al*., 2001).

The strong associations between gastro-intestinal helminths and gut bacteriota may be interpreted under two perspectives. First, the strong positive associations between the diversity of helminth community and gut bacteriota might corroborate the hypothesis. Besides, experimental evidence showed that helminths have the capacity to maintain higher gut microbiota diversity and may represent gut homoeostasis (Kreisinger *et al*., 2015). Indeed, low-intensity, chronic helminth infections are commonly linked to high microbial diversity and predominance of bacteria typically associated with gut health (Peachey *et al*., 2017).

Nevertheless, this interpretation has to be taken cautiously as the diversity of both communities was strongly influenced by the localities of sampling. The environment might therefore shape similarly gut bacteriota and helminth community diversity.

Second, significant associations between helminth community and gut bacteriota composition which remain significant even when potential geographic confounding effects were removed – may be linked to the fact that both communities reside in the same environmental niche (host intestines). From there, they likely experience similar selection pressure (e.g., host immune responses) which could shape their composition (Glendinning *et al*., 2014). One could expect therefore potentially strong interactions and reciprocal influence between them, Unfortunately, the causal processes behind these gut microbiota and helminths interactions are complex, multifaceted and difficult to assess. This intricacy is amplified by the fact that experimental studies mostly focus on single helminth infections while interactions between/within community are the rule within host organisms. The field of microbiota research would thus benefit from taking into account the whole composition of gastro intestinal helminth community rather than single helminth infections only.

In this study, we also highlight a large number of species-specific associations between helminths infections and members of the gut bacteriota. High-intensity, acute helminth infections may correlate with changes in hosts gut microbiota, through direct and indirect interactions (e.g., immune or other processes such as malnutrition; Peachey *et al*., 2017). Nevertheless, the patterns of shifts in gut bacteriota associated with helminth infections remain hardly predictable so far. As such, research works addressing this issue with laboratory or wild animals have provided variable, and sometimes even contradictory conclusions. Most surprising is that these inconsistent patterns are also found when focusing on single host-helminth models. A potential explanation is that these infection-associated microbiota shifts could depend on the presence of other helminths and the duration of infection (Sabey et al., 2021). Local interactions between helminths and between helminths and gut bacteria could mediate changes in infection outcomes as well as the gut bacteria and helminth populations themselves (Glendinning *et al*., 2014).

### Conclusion

Altogether, these results emphasize complex interlinkages between gut bacteriota, gastro-intestinal helminths and bacterial infections in wild animal populations. We emphasize the strong impact of environment, even at fine geographical scales, on these interactions. Shifts in diet or host genetics could mediate the spatial changes observed in gut bacteriota. However, the processes shaping gut bacteriota diversity and composition are many and complex, and further investigations are required to decipher the relative importance of drift, dispersal or selection on bank vole gut bacteriota in the populations studied here. Besides, we find a diverse array of associations between gut bacteriota and gastro-intestinal helminths or pathogenic bacteria, some being significant at the scale of the whole community and other being species-specific only. Whether these patterns reflect coadaptation, dysbiosis or indirect interactions with host immunity and coinfections should now be considered to better understand the spatial variations observed in the relationships between gut bacteriota and health.

## Acknowledgement

Data used in this work were partly produced through the genotyping and sequencing facilities of ISEM (Institut des Sciences de l’Evolution-Montpellier) and Labex CeMEB (Centre Méditerranéen Environnement Biodiversité).

## Data and script availability

Raw data and scripts are available on zenodo: https://doi.org/10.5281/zenodo.7433800

## Supplementary information

All supplementary materials are available on zenodo : https://doi.org/10.5281/zenodo.7431842

**Supplementary Figure S1**. Maps showing the sampling area (left) and localities (right) in France. Forests are indicated in green and water in blue. The four sampling localities are represented with a colored polygon. The arrow indicates the North.

**Supplementary Figure S2**. Composition of the gut bacteriota. The relative abundance of six phyla representing 99% of the total composition is represented. Individuals are grouped by sampling localities, which are ordered from North to South. (A) Bar graph shows individual variation in phyla composition (phylum=color). (B) Box and whisker plots represent median and interquartile values for each phylum. Black dots correspond to mean values, and colored dots correspond to individuals.

**Supplementary Figure S3**. Variations of alpha diversity with individual factors, for the gut bacteriota (family level), pathogenic bacteria and gastro-intestinal helminths of bank voles. Alpha diversity is estimated using the specific richness (A, B and C) and the Shannon index (D, E and F). In graphs C and F, the blue line corresponds to the linear regression line.

**Supplementary Figure S4**. Relationships between the composition of the gut bacteriota, pathogenic bacteria and gastro-intestinal helminth communities: The db-RDA triplot shows the structure of the gut bacteriota at the phylum level and the correlations with the intra-host parasite communities. The arrows correspond to the significant explanatory variables. Each point corresponds to an individual, and the colors correspond to the different sampling localities.

**Supplementary Table S1**. Variation of the Firmicutes/Bacteroidetes ratio with localities and individual factors.

**Supplementary Table S2**. Alpha diversity metrics and statistics for the gut bacteriota, pathogenic bacteria and helminth communities of bank voles.

**Supplementary Table S3**. Beta diversity metrics and statistics for the gut bacteriota, pathogenic bacteria and helminth communities of bank voles.

## AUTHOR CONTRIBUTIONS

M.B.: Data curation; Formal analysis; Methodology; Writing – original draft

M.G.: Conceptualization; Data curation; Formal analysis; Methodology; Supervision; Writing review and editing

A.D.: Conceptualization; Data curation

C.A.D.: Data curation; Writing – review and editing

P.M.: Conceptualization; Funding acquisition; Investigation; Supervision; Writing – review and editing

B.R.: Conceptualization; Methodology; Supervision; Writing – review and editing

N.C.: Conceptualization; Funding acquisition; Investigation; Methodology; Project administration; Resources; Supervision; Writing – original draft

