## Supplementary Figures for "Three-way relationships between gut microbiota, helminth assemblages and bacterial infections in wild rodent populations"

**Figure S1. Map showing the sampling area (left) and localities (right) in France.** Forests are indicated in green and water in blue. The four sampling localities are represented with a colored polygon. The arrow indicates the North.

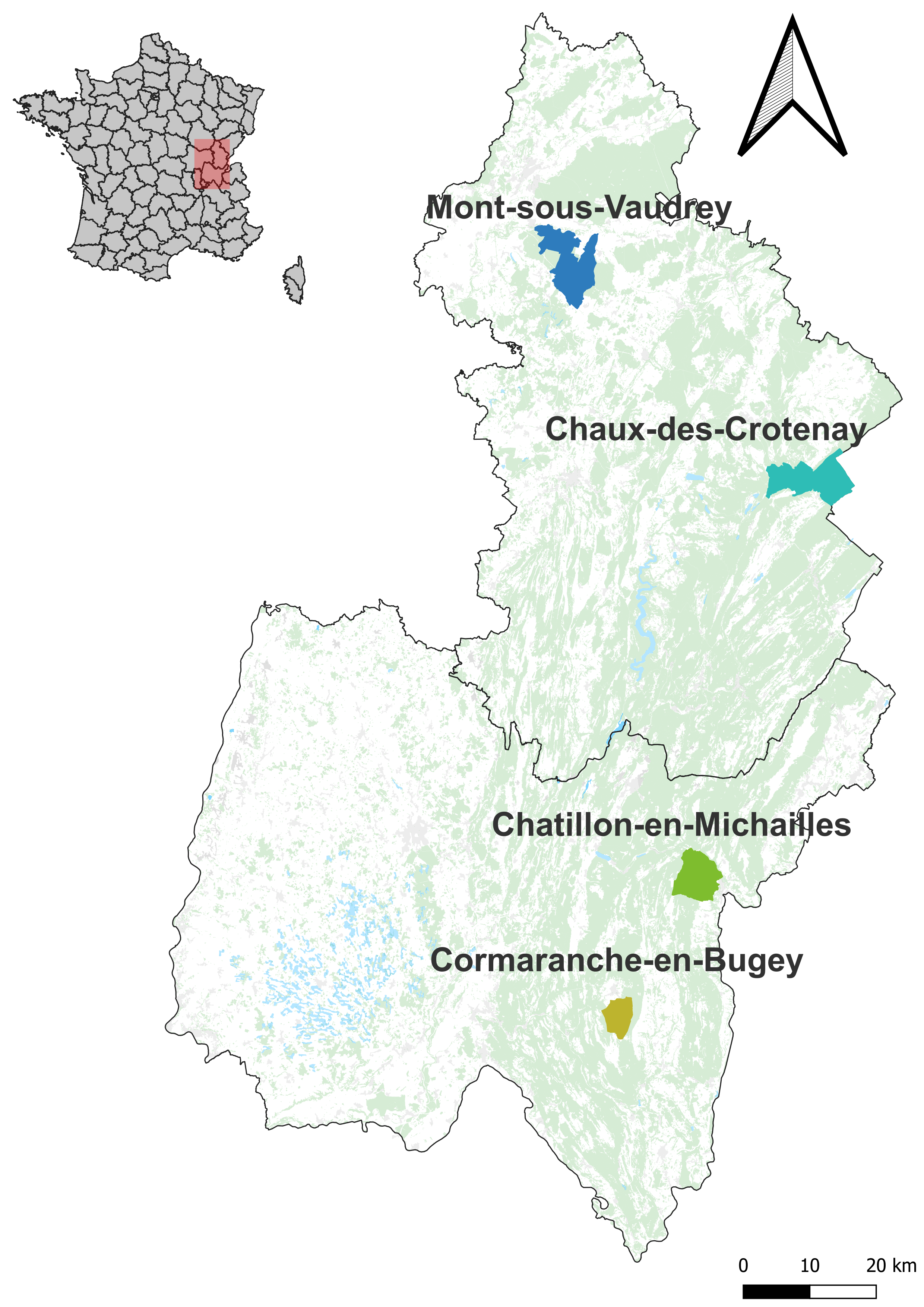

**Figure S2. Composition of the gut bacteriota of *M. glareolus*.** The relative abundance of six phyla representing 99% of the total composition is represented. Individuals are grouped by sampling localities, which are ordered from North to South. (A) Bar graph shows individual variation in phyla composition (phylum=color). (B) Box and whisker plots represent median and interquartile values for each phylum. Black dots correspond to mean values, and colored dots correspond to individuals.

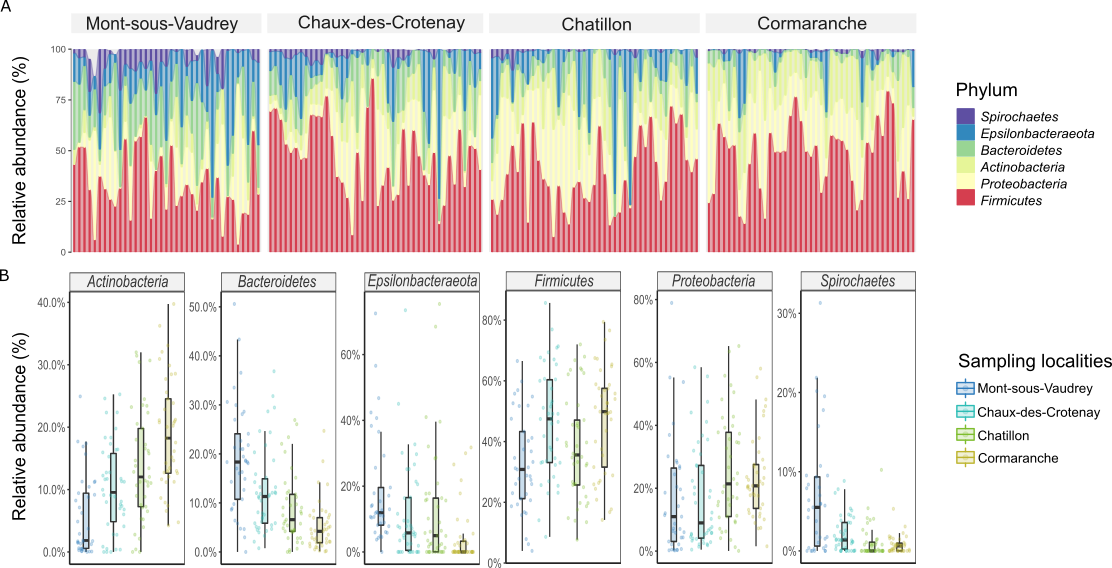

**Figure S3.** **Variations of alpha diversity with individual factors, for the gut bacteriota (family level) of bank voles.** Alpha diversity is estimated using the specific richness (A, B and C) and the Shannon index (D, E and F). In graphs C and F, the blue line corresponds to the linear regression line.

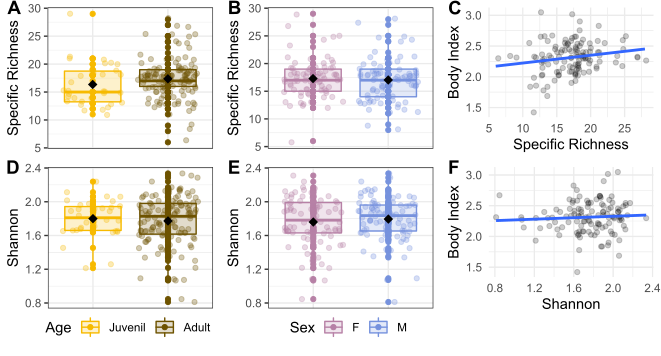

**Figure S4.** **Relationships between the composition of the gut bacteriota, pathogenic bacteria and gastro-intestinal helminth communities:** The db-RDA triplot shows the structure of the gut bacteriota at the phylum level and the correlations with the intra-host parasite communities. The arrows correspond to the significant explanatory variables. Each point corresponds to an individual, and the colors correspond to the different sampling localities.

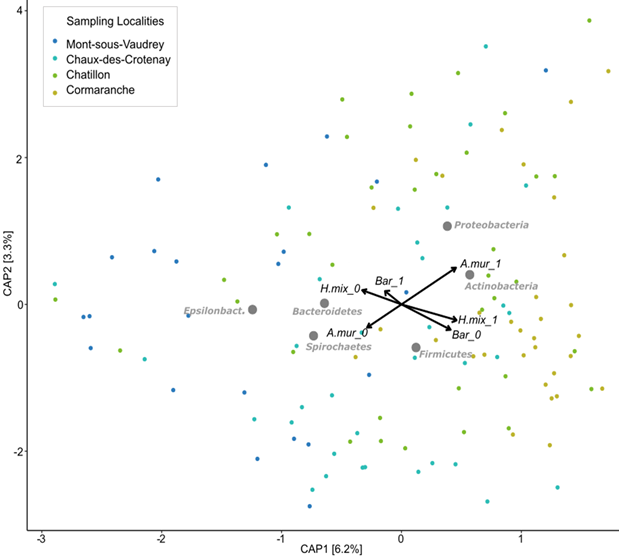
